# STARD3 mediates non-vesicular cholesterol transport in *Caenorhabditis elegans*

**DOI:** 10.64898/2026.01.09.698688

**Authors:** Bernabe Battista, Agustina Sambrailo, Franco A. Biglione, Verónica A. Lombardo, María C. Mansilla, Daniela Albanesi, María-Natalia Lisa, Diego de Mendoza, Andres Binolfi

## Abstract

Cholesterol transport plays a pivotal role in regulating development and metabolism in *Caenorhabditis elegans*, a sterol-auxotrophic organism. Here, we identify the nematode cholesterol-binding protein STARD3 and provide structural and functional evidence for its role in non-vesicular sterol mobilization. Using biophysical and high-resolution structural methods, we show that the START domain (Ce-START) of *C. elegans* STARD3 binds cholesterol with high affinity and adopts a fold conserved with its human ortholog. Crystal structures of Ce-START in both apo and cholesterol-bound forms reveal key determinants of sterol recognition and conformational changes upon ligand binding. Functional analysis of a *C. elegans stard3* knockout strain demonstrates that STARD3 is essential for cholesterol trafficking under sterol-limited conditions and that it genetically interacts with the NPC1/NPC2 pathway to sustain cholesterol mobilization. Collectively, these results establish STARD3 as a crucial cholesterol transporter in *C. elegans* and underscore the evolutionary conservation of START-domain proteins, reinforcing the utility of *C. elegans* as a model for studying intracellular cholesterol dynamics.

## Introduction

Cholesterol is a structural component of cell membranes and a precursor of steroid hormones and bile acids and is therefore essential for development and health (1). Both cholesterol excess and cholesterol deficiency can have detrimental effects on health, and a myriad of regulatory processes have thus evolved to control the metabolic pathways of sterol metabolism (2). Much of our knowledge of these systems, their rate-controlling processes, and how they mechanistically affect health and life span comes from studies on the nematode *Caenorhabditis elegans* (3,4). *C. elegans* are auxotrophic for sterols: in contrast to mammals, they cannot synthesize cholesterol, and dietary supply is therefore essential for survival. In *C. elegans*, cholesterol regulates at least two processes. First, it is required for growth and progression through larval stages, as well as for proper shedding of old cuticles during the molting events (5). Second, it regulates entry into a specialized diapause stage for survival under unfavorable conditions, called the dauer larva (6). Tight spatial and temporal regulation of uptake, storage, and transport of sterols to appropriate subcellular compartments is stringently required for cholesterol to exert its diverse cellular functions. The sterols that govern dauer formation are bile-acid-like hormones called dafachronic acids (DAs). Molecular mechanisms underlying their function have been intensely studied. DAs inhibit dauer formation by binding to the nuclear hormone receptor DAF-12, which, in the absence of DAs, activates the dauer program (7,8).

Following cholesterol intake, its systemic distribution in worms is mediated by various transport systems, including the vitellogenin lipoprotein particles (9). After receptor-mediated endocytosis of cholesterol carriers, cholesterol is trafficked through the endo-lysosomal system and directed to other cellular compartments with the assistance of the Niemann-Pick type C1 (NPC1) homologs NCR-1 and NCR-2 (10). The importance of this process is demonstrated by the fact that *ncr-2-*(*-*)*;ncr-1-*(*-*) null mutants fail to produce fertile adults and instead arrest at the dauer diapause(11). This developmental arrest occurs as a result of a deficiency in cholesterol trafficking and decreased DA production (12).

During the last decade, studies on mammalian cells showed that carriers such as the steroidogenic acute regulatory proteins (StAR) provide an important pathway for the transfer of cholesterol from the endosomal-lysosomal system to other cellular organelles at membrane contact sites (13). STARD3 is a member of the StAR protein family that binds cholesterol. It is ubiquitously expressed and anchored to the membrane of late endosomes/lysosomes (LE/Ly) (14). Its function relies on two key features: its ability to establish contact sites between LE/Ly and the endoplasmic reticulum (ER) through interaction with VAP proteins, and its capacity to exchange sterols via its START domain (13). The protein STARD3 regulates intracellular cholesterol distribution and organelle dynamics, supporting lipid homeostasis, particularly under stress or pathological conditions such as Niemann–Pick C disease (15,16). STARD3 expression has been associated with increased cholesterol content in various organelles, including endosomes, mitochondria, and the plasma membrane (17).

Structurally, mammalian STARD3 comprises two functional divergent domains known as MENTAL and START, connected by a flexible linker harboring a FFAT (two phenylalanines in an acidic tract) recognition motif (18). While MENTAL is responsible for STARD3 anchoring to LE/Ly membranes through its transmembrane helices (17), the cytosolic START domain contains a conserved hydrophobic cavity that binds cholesterol with high affinity (13,19). In that sense, the START domain functions as a “hydrophobic bridge” across the aqueous compartments between membranes (20). Despite these evidence, the molecular details of the role of STARD3 in cholesterol metabolism are still scarce, in part due to the absence of high-resolution tridimensional structural data of cholesterol:START complexes and the lack of simple animal models to perform biochemical, genetic and functional studies of STARD3-cholesterol interactions.

In this work, we conducted a structural and functional characterization of the molecular features driving cholesterol recognition in the *C. elegans* STARD3 homolog (Ce-STARD3). Using a variety of low- and high-resolution biophysical methodologies we determined that the START domain of *C. elegans* STARD3 (Ce-START) binds cholesterol with high affinity and a 1:1 stoichiometry. The structure of the START-cholesterol complex allowed us to delineate the molecular interactions taking place within the lipid-binding cavity, providing unique insights into the structural determinants responsible for cholesterol recognition and complex stabilization in STARD3. Next, we constructed a *C. elegans* strain where the *stard3* gene was knocked out and performed functional and biochemical assays, disclosing a role for STARD3 in non-vesicular cholesterol mobilization that depends on the interactions with the NPC1 and NPC2 system. Overall, our results show that, like its human ortholog, STARD3 functions as a lipid transfer protein and is involved in cholesterol flux in *C. elegans*. These findings further support the use of this animal model to understand the role of cholesterol metabolism in both physiological and pathological processes.

## Material and Methods

### Reagents

Cholesterol, water-soluble cholesterol (ChlCDX) and methyl-β-cyclodextrin (mCDX) were purchased from Sigma (Sigma-Aldrich, St. Louis, Missouri, USA).

### Cloning, expression and purification of Ce-START

The sequence for *C. elegans* STARD3 domain was identified in the worm base (WormBase) using a template from the human ortholog (Uniprot Q14849). The *C. elegans* gene is annotated as TAG-340 (WormBase) and it is predicted to be an ortholog of human STARD3 (21). We selected the sequence corresponding to the START domain, synthesized the gene adding a 6xHis tag and a TEV cleavage site (Tobacco etch protease) to the N-terminus and then cloned into pET15b vector (General Biosystems). The START construct was overexpressed in the BL-21 (λDE3) C41 cells at 20°C using 0.5 M IPTG (isopropyl β-D-thiogalactoside) for 16 h. The bacterial pellets were lysed in 300 mM NaCl, 50 mM Na_2_HPO_4_ (pH 8.0), 1mM PMSF (phenylmethylsulphonyl fluoride), followed by centrifugation for 30 minutes at 20000 *g*. Enrichment of the protein from the supernatant was carried out using a nickel-affinity chromatography according to the manufacturer’s protocol (GE healthcare). Protein was eluted in a buffer consisting of 300 mM NaCl, 50 mM Na_2_HPO_4_ (pH 8.0) and with a 10 to 1000 mM imidazole gradient. We then exchanged this buffer for another one consisting of 150 mM NaCl, 20 mM Na_2_HPO_4_ (pH 7.0), 1mM DTT (Dithiothreitol) using a standard dialysis membrane tubbing of MWCO 3.5 kDa (SpectraPor) in the presence of TEV protease at a ratio of 1:100. We checked the integrity of the protein and the complete removal of the 6xHis by SDS-PAGE analysis. The protein was concentrated using Amicon ultracentrifugation filters (Merck) and further purified employing an ÄKTA Pure FPLC instrument (Cytiva) and a size exclusion column Superdex^®^75 10/300 (Amersham Biosciences). The elution was carried out using a buffer consisting of 20 mM Na_2_HPO_4_ (pH 7), 150 mM NaCl and 1mM DTT for NMR/CD/Fluorescence experiments and 25 mM Tris-HCl (pH 7.0), 150 mM NaCl and 1mM DTT for the crystallization assays. The elution fractions corresponding to the maximum of the 280 nm absorption band were collected, and the accurate molecular weight (26.1 kDa) was verified by SDS-PAGE analysis. The final Ce-START concentration was estimated by absorbance at 280 nm, considering a molar absorption coefficient of 35000 M^−1^cm^−1^.

### Sequence alignment and structural modelling of Ce-STARD3

Sequence alignment was performed using ClustalW on the ESPript3 platform with default parameters (22). The sequences employed were UniProt Q14849 (human STARD3) and UniProt Q19819 (Ce-STARD3). The structural modelling was conducted using ColabFold v1.5.5: AlphaFold2 (MMseqs2) (23) and the images were generated with PyMOL (The PyMOL Molecular Graphics System, Schrödinger, LLC)

### CD spectroscopy

CD measurements were conducted on a Jasco J-1500 spectrometer with a 1 mm path-length quartz cuvette and a protein concentration of 8 µM. The effect of cholesterol binding on the protein’s secondary structure was assessed by monitoring changes in ellipticity within the far-UV region (190–260 nm) upon lipid titration. Thermal denaturation curves of the protein in the absence or presence of cholesterol were obtain by monitoring CD values at 222 nm as the temperature increased from 15°C to 81°C. Prior to the denaturation experiments Ce-START was incubated with a 4-fold excess of cholesterol (ChlCDX) for 30 minutes at 20°C to ensure full complex formation. Experiments were repeated at least two times. In all cases, control experiments using cholesterol-free mCDX were collected. The temperature-induced denaturation curves were obtained following previous studies with other START domains (24,25).

### Intrinsic fluorescence measurements

Intrinsic tryptophan fluorescence measurements were performed using a Cary Eclipse Fluorimeter at 20°C. The instrument was configured with an excitation wavelength of 280 nm and an emission wavelength range of 300 to 500 nm, using a 5 nm slit width for excitation and emission. Increasing concentrations of Chl:mCDX or mCDX samples (prepared in the same buffer as the protein) were added to a 10 μM Ce-START solution. After incubating the samples for 30 minutes to allow the system to reach equilibrium, fluorescence measurements were recorded. To determine the specific contribution of cholesterol binding to fluorescence quenching, the mCDX contribution was subtracted from each added concentration, resulting in difference fluorescence spectra. Complex formation was quantified by the decreased in initial protein fluorescence, expressed as fraction quenching (ΔF/F_0_), where F_0_ and ΔF correspond to the integrated fluorescence intensities over the 320-360 nm range for the initial sample (without Chl) and for the absolute difference spectra of each assayed cholesterol concentration, respectively. Fluorescence integrals were calculated using Riemann’s summation. Determinations were performed as triplicates.

### NMR spectroscopy

The NMR experiments were conducted using a 700 MHz Avance III spectrometer, equipped with a 5 mm TXI inverse detection probe (^1^H/D-^13^C/^15^N) with magnetic field gradients along the z-axis. All spectra were acquired and processed using the Topspin 3.5 software (Bruker, BioSpin). The ^1^H-^15^N HSQC experiments (*fhsqcf3gpph*, Topspin library) were performed at 25 °C on 150 µM ^15^N uniformly-enriched Ce-START samples. A total of 2048 and 256 points were acquired for ^1^H and ^15^N, respectively, for sweep widths of 15.9 and 40 p.p.m. We used a relaxation delay of 1 s and 64 scans. The spectra were processed using zero filling (two times the number of acquired points), apodization by multiplication with a sine function in both dimensions, Fourier transform, and baseline correction.

### Ce-START crystallization, data collection and structure determination

Crystallization trials of native Ce-START (2 mM), alone or in the presence of cholesterol (1.5 mM) were carried out using the vapor diffusion method, employing a Gryphon nanoliter-dispensing crystallization robot (Art Robbins Instruments), by mixing 250 nL of protein solution and 250 nL of reservoir solution, equilibrated against 80 µl of reservoir solution, in 96-well Intelli-Plates (Hampton). Crystallization plates were stored at 20°C and inspected with a stereo microscope (Leica) to monitor crystal growth. The optimized conditions for crystal growth were 100 mM sodium citrate tribasic dihydrate pH 5.0, 200 mM lithium sulfate; 22% v/v PEG 200 for Ce-START crystals and 50 mM sodium cacodylate pH 6.5, 100 mM magnesium acetate, 200 mM potassium chloride, 14% w/v PEG 8000 for Ce-START plus cholesterol (Ce-START:Chl) crystals. Ce-START and Ce-START:Chl crystals grew to a size of ca. (50 µm)^3^ in both cases. Crystals were flash-frozen in liquid nitrogen, using 25% v/v glycerol as cryoprotectant, for data collection at 100 K.

X-ray diffraction data were collected from Ce-START and Ce-START:Chl single crystals on beamline I03 (Diamond Light Source, UK) using wavelength 0.976230 Å. Diffraction data were indexed and integrated with XDS (26) and scaled with Aimless (27) from the CCP4 program suite (28). Ce-START and Ce-START:Chl crystals belonged to the space group P2_1_, although they were not isomorphous, and diffracted to high resolution. Both crystal structures were solved by molecular replacement with Phaser (29) using a Ce-START model spanning residues 213-447 of the protein built by Alpha Fold 2(23) as a search probe. In the case of the structure of Ce-START:Chl, cholesterol molecules were manually placed in a *mFo–DFc* electron density map obtained after successive cycles of manual model building with Coot (30) and crystallographic refinement with phenix.refine (31). For the two structures presented in this work, atomic coordinates and individual B-factors were refined to convergence with phenix.refine (31) using torsion-angle NCS restraints and TLS parameters. Final models were validated through the Molprobity server (32).

### C. elegans strains

The worm strains Bristol N2 and *ncr-2-*(*-*)*; ncr-1-*(*-*) (JT10800) were obtained from Caenorhabditis Genetics Center (CGC, University of Minnesota). The *stard3-*(*-*) knockout strain PHX3367 [*tag-340*(syb3367)] was purchased from Sunny Biotech (China). It was generated by deleting 2204 bp from the *stard3-*(*-*) gene (*tag-340*). The deletion was confirmed by PCR analysis using the following primers (5’-3’): FWD Out, tgatatcagttgtctcattcgtg; REV Out, agaaagggcaaggaggtatg and REV Int, ctctggattctcgattccg.

### Growth and maintenance of worm strains

Worms were routinely handled following previously established routines (33). Briefly, worms were propagated on nematode growth medium (NGM) agar plates seeded with *E. coli* OP50 (3). Saturated cultures of *E. coli* OP50 grown in lysogeny broth (LB) medium were concentrated 10 times and spread on NGM-agar plates. *ncr-2-*(*-*)*; ncr-1-*(*-*) mutants were grown at 15 °C and then shifted to 20°C to exacerbate the dauer larvae phenotype.

### Preparation of sterol-depleted plates and sterol-deprived worm culture

To obtain sterol-free conditions, agar was replaced by ultrapure agarose and peptone was omitted from plates (33). Briefly, agarose was washed three times overnight with chloroform to deplete trace sterols. Salt composition was kept identical to NGM plates. As a food source, *E. coli* NA22 grown overnight in sterol-free culture medium DMEM was used. Bacteria were rinsed with M9 buffer and concentrated 20 times. Bleached embryos were grown for one generation on sterol-free agarose plates. The resulting gravid adults were bleached, and the obtained embryos were used in various assays. The animals grown for two generations in the absence of cholesterol arrest as L1-L2* larvae (34). The bypass of the L2* larval arrest is indicated as % of developed adults.

### RNAi by feeding

RNAi against *stard3* was performed with *E. coli* HT115 double-stranded RNA (dsRNA)-producing strain from the Ahringer library (35,36). RNAi was performed in the first generation. Bleached embryos were left to hatch overnight in M9 at 20°C and the resulting L1s were seeded to RNAi plates and grown at 20°C. Finally, the dauer phenotype was scored after 120 h (37).

### Scoring brood size, embryonic lethality, and larval lethality

To assess brood size, embryonic lethality, and larval lethality, age-matched L4 hermaphrodites, grown on cholesterol-free medium or standard NGM plates, were transferred daily to fresh plates of the same composition for four consecutive days. The total number of fertilized eggs laid, hatched, and the number of progenies that reached adulthood were scored. At least 10 worms per condition were analyzed. Larval lethality was not quantified under cholesterol-free conditions, as the second-generation progeny (F2) grown in the absence of cholesterol arrest at the L2* or dauer-like stage (34).

### Dauer formation assays

In general, 50–80 L1s were transferred to NGM plates seeded with *E. coli* (HT115) containing RNAi against *stard3* or empty vector as a negative control. The supplementation with cholesterol 1X or 10X (final concentration 13 µM and 130 µM, respectively) was added to the NGM media according to the volume used for the preparation of the plates. Finally, after 120 h the percentage of dauers was scored (33).

## Results

### Characterization of the cholesterol binding features of *C. elegans* STARD3

To understand the role of the STARD3 protein in the mobilization of internal cholesterol reserves in *C. elegans*, we used the WormBase database to search for a gene coding for STARD3 protein. We found that the gene annotated as *tag-340* codifies for a putative STARD3 ortholog (UniProt accession number Q19819) (21). Despite only sharing 28.68% of identity with the human STARD3 sequence (UniProt Q14849, **Supplementary Fig. 1**), structural alignment of TAG-340 AlphaFold2 model with the corresponding human model revealed a high degree of similarity, suggesting a conserved function in intracellular cholesterol transport within the two species (**Supplementary Fig. 2 and 3**).

To clarify its function, we conducted a biophysical characterization of the interaction between the Ce-START domain and cholesterol. We recombinantly purified ^15^N isotopically-enriched Ce-START protein (**Supplementary Fig. 4A**) and recorded ^1^H-^15^N HSQC spectra in the absence and presence of cholesterol (**Fig. 1A, B**). The splitting of signals at a 0.75:1 cholesterol:Ce-START ratio, which converged to the new detected species when we added an excess of 2 cholesterol equivalents, provided evidence for cholesterol interaction with this domain. We did not detect concentration-dependent chemical shifts perturbations which are indicative of a fast exchange regime of cholesterol at the binding site (38). Instead, the disappearance of signals from the free protein state and the build-up of new signals in the presence of cholesterol reflect a slow exchange binding regime (39). Thus, the spectral changes resemble two states of the protein, the free form and the cholesterol-bound form. This implies that the added cholesterol binds to the protein with a significantly lower dissociation constant than the protein concentration used in the experiment (150 µM) (39).

**Figure 1.**
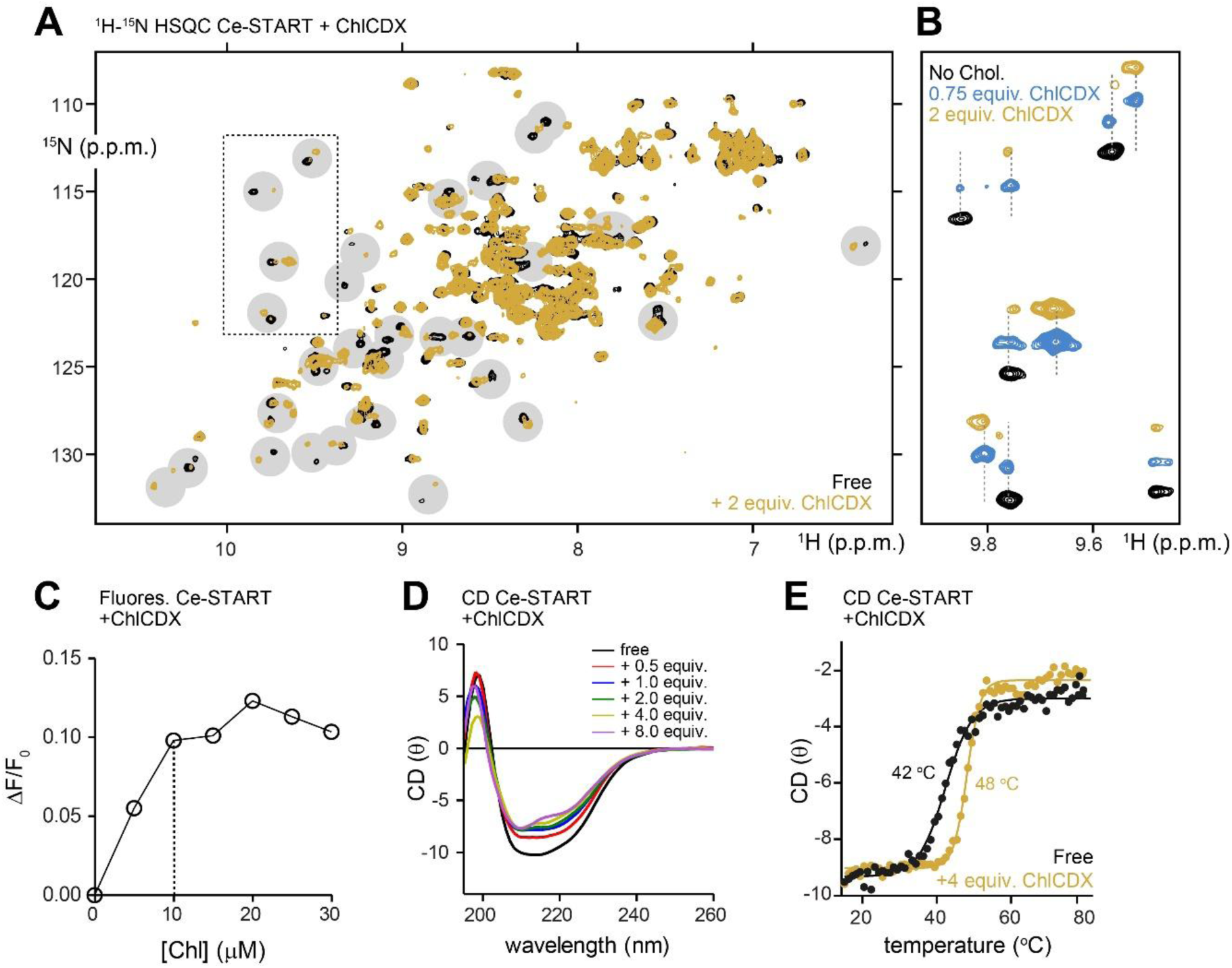
Biophysical characterization of cholesterol binding to Ce-START. **(A)** Overlay of ^1^H-^15^N HSQC spectra of 150 µM, ^15^N-isotopically enriched Ce-START in the absence (black) and presence (yellow) of 2 equivalents of cholesterol, added as a methyl-cyclodextrin complex (ChlCDX). The mixture was incubated at room temperature for 30 min to reach equilibrium before data acquisition. Cross-peaks exhibiting changes upon cholesterol addition are indicated with gray circles. **(B)** Enlarged view of the dashed-line box in panel **(A)**, showing signal doubling upon the addition of 0.75 equivalents (blue) or 2 equivalents (yellow) of cholesterol to Ce-START. NMR spectra were artificially shifted in ^15^N (vertical axis) for better visualization of cross-peaks corresponding to the apo or cholesterol-bound protein states. **(C)** Tryptophan fluorescence quenching of 10 µM Ce-START in the presence of increasing ChlCDX concentrations. Curve shows the fraction fluorescence quenching produced by Chl binding to Ce-START. **(D)** Far-UV CD spectra of 8 µM Ce-START in the absence (black line) or presence of increasing concentrations of ChlCDX (colored lines). **(E)** Ellipticity curves [θ] at 222 nm obtained from thermal denaturation of 8 µM Ce-START in the absence (black) and presence of 4 cholesterol equivalents (red).

Since cholesterol is insoluble in aqueous buffer, we used it in a complex with soluble methyl-β-cyclodextrin (mCDX). To confirm that the perturbations observed in **Fig. 1A, B** were caused by sterol binding and not by non-specific effects of mCDX, we performed a control experiment by adding the same proportions of free mCDX to Ce-START (**Supplementary Fig. 4B**). We did not detect any changes in the intensity or the chemical shift of the signals, confirming that the perturbations observed in Ce-START are due to cholesterol binding. With these data, we can also conclude that Ce-START can efficiently remove cholesterol from the ChlCDX complex. The affinity constant (*Ka*) of the ChlCDX complex is 1.7 × 10^4^ M^−1^ (*Kd* 59 µM) (40), indicating that the affinity of the interaction between Ce-START and cholesterol is higher than that.

Next, we used fluorescence quenching to further analyze the binding process. We titrated cholesterol into a 10 µM solution of Ce-START and followed the fluorescence intensity of tryptophan (Trp) sidechains (41). After subtracting the unspecific effects of mCDX (**Supplementary Fig. 4C-F**), we observed a quenching effect on the Trp fluorescence that was dependent on the concentration of added cholesterol (**Fig. 1C**), confirming that Ce-START binds to cholesterol. However, it was not possible to determine the precise value of the dissociation constant, as it is lower than the protein concentration used. Thus, the binding curve accurately reflects the stoichiometry but not the affinity (42). In this way, we were able to determine that each molecule of Ce-START binds one molecule of cholesterol and that the *Kd* of the complex is lower than 10 µM. These results agree with previous evidence reporting a 1:1 interaction and a *Kd* of approximately 30 nM between a human START domain and cholesterol (25).

Although initial NMR analysis revealed that ligand binding induces changes in Ce-START, the lack of residue-specific assignment prevented us from determining whether conformational changes are associated with the interaction. Thus, we complemented the biophysical analysis using circular dichroism (CD) spectroscopy. Upon addition of increasing amounts of cholesterol, we detected concentration dependent changes in the CD spectra (**Fig. 1D**). In particular, we observed that the maximum at 199 nm and the minimum at 214 nm were attenuated and that this effect exclusively depended on the interaction between cholesterol and Ce-START (**Supplementary Fig 4G, H**). These results suggest that cholesterol binding elicits a minor decrease in α-helical secondary structure content (43). We also evaluated the conformational stability of the Ce-START in the absence and presence of cholesterol using CD spectroscopy at increasing temperatures (43). As shown in **Fig. 1E**, the denaturation profile of Ce-START exhibited a single, cooperative transition, pointing to a well-defined tertiary structure and a two-state unfolding process. This behavior is similar to that observed in other START domains (24,25). The unbound protein exhibited a melting temperature (Tm) of 42°C, which increased to 48°C in the presence of cholesterol, indicating that lipid binding stabilizes the Ce-START overall structure.

### Structure of the Ce-START:Chl complex

To analyze the binding of cholesterol to Ce-START with higher resolution we solved crystal structures of Ce-START in the absence and presence of cholesterol (**Table 1**). The crystal structure of Ce-START without ligand contained two molecules per asymmetric unit, whereas the cholesterol-bound contained four. In both cases, electron density maps provided evidence for the region spanning residues 233-443, although depending on the protein chain it was possible to refine some additional residues at the disordered N- and C-termini and/or short internal loop segments. Superposition by secondary structure matching displayed an average RMSD of 0.5 Å and never exceeded 0.9 Å in whole-chain comparisons performed for either or both structures. This indicates that the overall fold of Ce-START is independent of crystal packing and ligand binding.

**Table 1.**
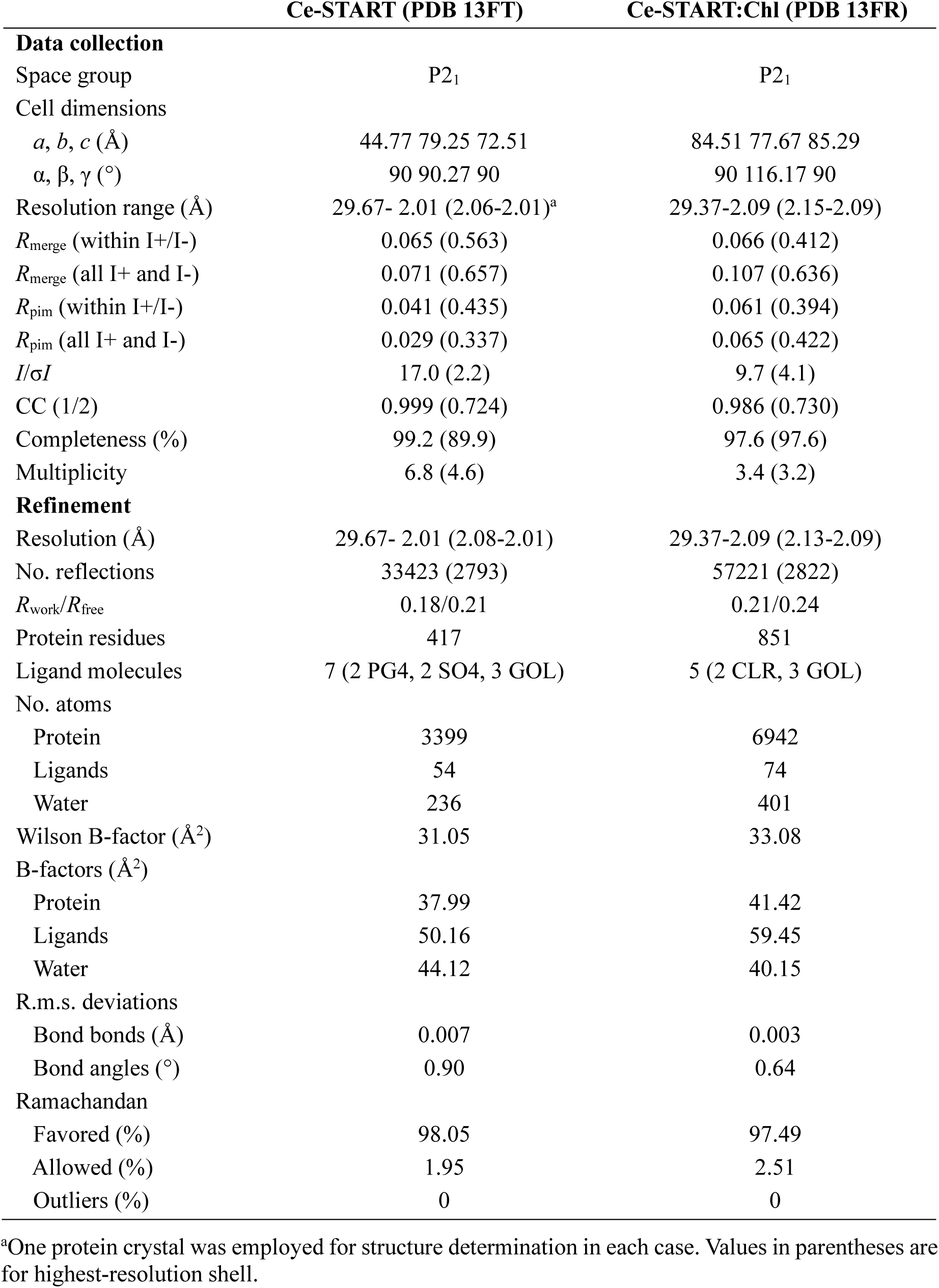
X-ray diffraction data collection and refinement statistics.^a^.

The tertiary structure of Ce-START closely resembles the one of the START domain from human STARD3 (20,44) and other STARD proteins (13,19,45,46) (**Supplementary Fig. 5A, B**). It consists of a nine-stranded twisted antiparallel β-sheet and three α-helices packed such that a cholesterol binding groove is formed on the concave side of the β-sheet array (**Fig 2A**). These antiparallel β-sheet form an incomplete β-barrel with a U-shape while alpha-helices α2 and α3 serve as the cavity roof.

**Figure 2.**
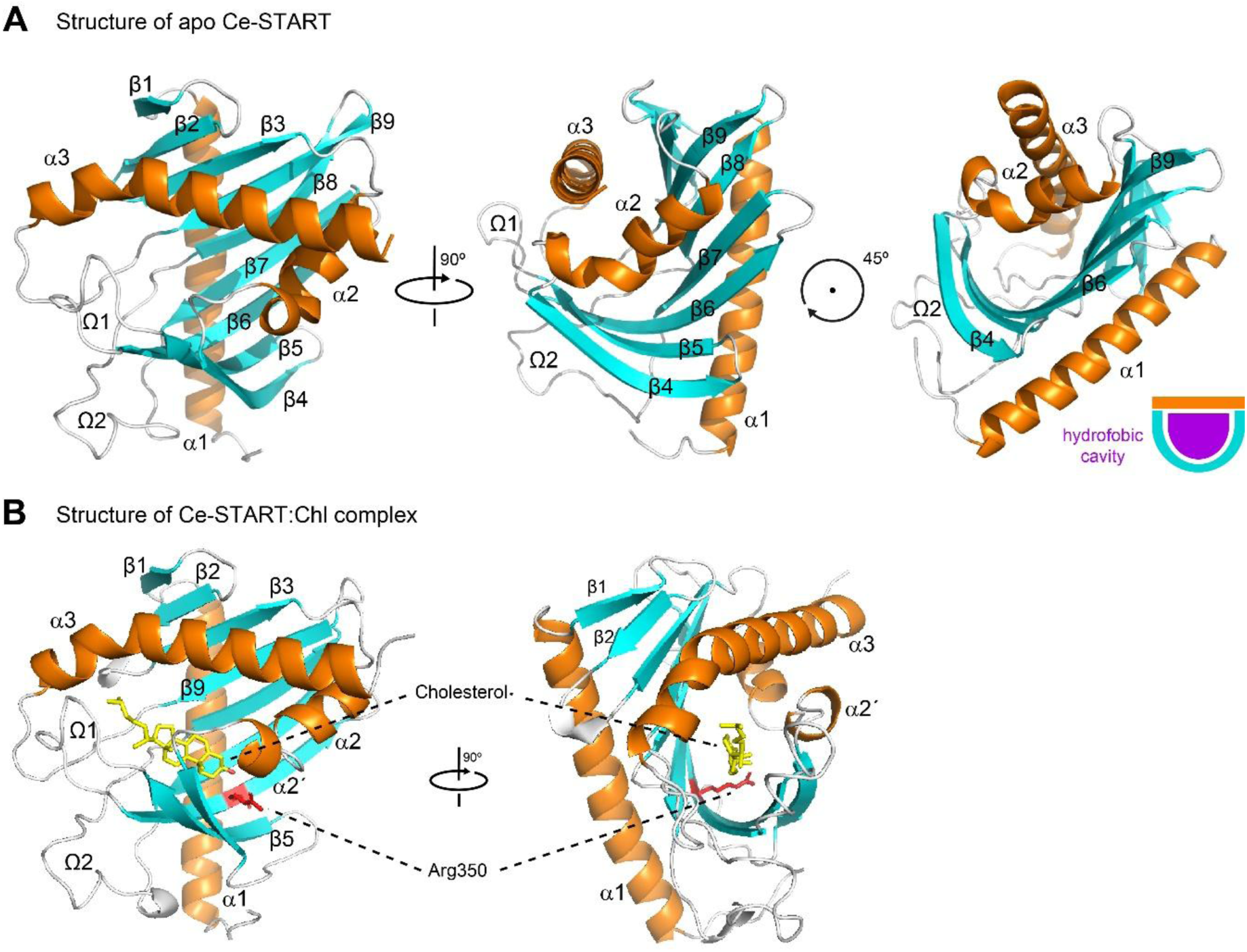
Structure of Ce-START domains. **(A)** Crystal structure of the Ce-START domain. The left panel shows a side view of the domain, oriented perpendicular to the hydrophobic cavity. The central panel presents a top view of the cavity, while the right panel displays the same perspective rotated 45° relative to the central panel, allowing visualization of one of the open entrances to the hydrophobic cavity. The diagram in the lower right illustrates the composition of the cavity: its walls and base are formed by an incomplete U-shaped β-barrel, while its roof is defined by helices α2 and α3 along with the Ω1 loop. **(B)** Crystal structure of the Ce-START domain in complex with cholesterol. The left and right panels show side and top views of the hydrophobic cavity, respectively. Arginine residue 350 is highlighted in red, while the cholesterol molecule (yellow) is positioned inside the cavity. The hydroxyl group of cholesterol is indicated in red.

To date, no experimental structures of START proteins complexed with cholesterol have been reported, and insights into cholesterol-binding features of START domains have relied primarily on molecular docking studies (20). Thus, our results provide experimental evidence of cholesterol binding to a START domain. In the Ce-START structure, cholesterol is bound with the α-face of the steroid nucleus packed against the proteińs β-sheet, while its hydroxyl group forms a hydrogen bond with Arg350 (**Fig. 2B**). Cholesterol binding alters the torsion angle of helix α3, increasing from 18.2° in Ce-START to 22.1° in Ce-START:Chl and the distances between loop Ω1 (Ala336) and the nearest residues in helix α3 (Gly423 and Tyr427) decreased from 9.7 Å and 11.6 Å to 8.4 Å and 9.7 Å, respectively (**Fig. 3A**). In addition, we detected a ligand-induced rearrangement involving the fragmentation of helix α2 into two shorter helices, α2 and α2′ (**Fig. 3A**). These structural perturbations are consistent with the decrease of ellipticity at 222 nm observed in the CD spectra of Ce-START bound to cholesterol (**Fig. 1D**).

**Figure 3.**
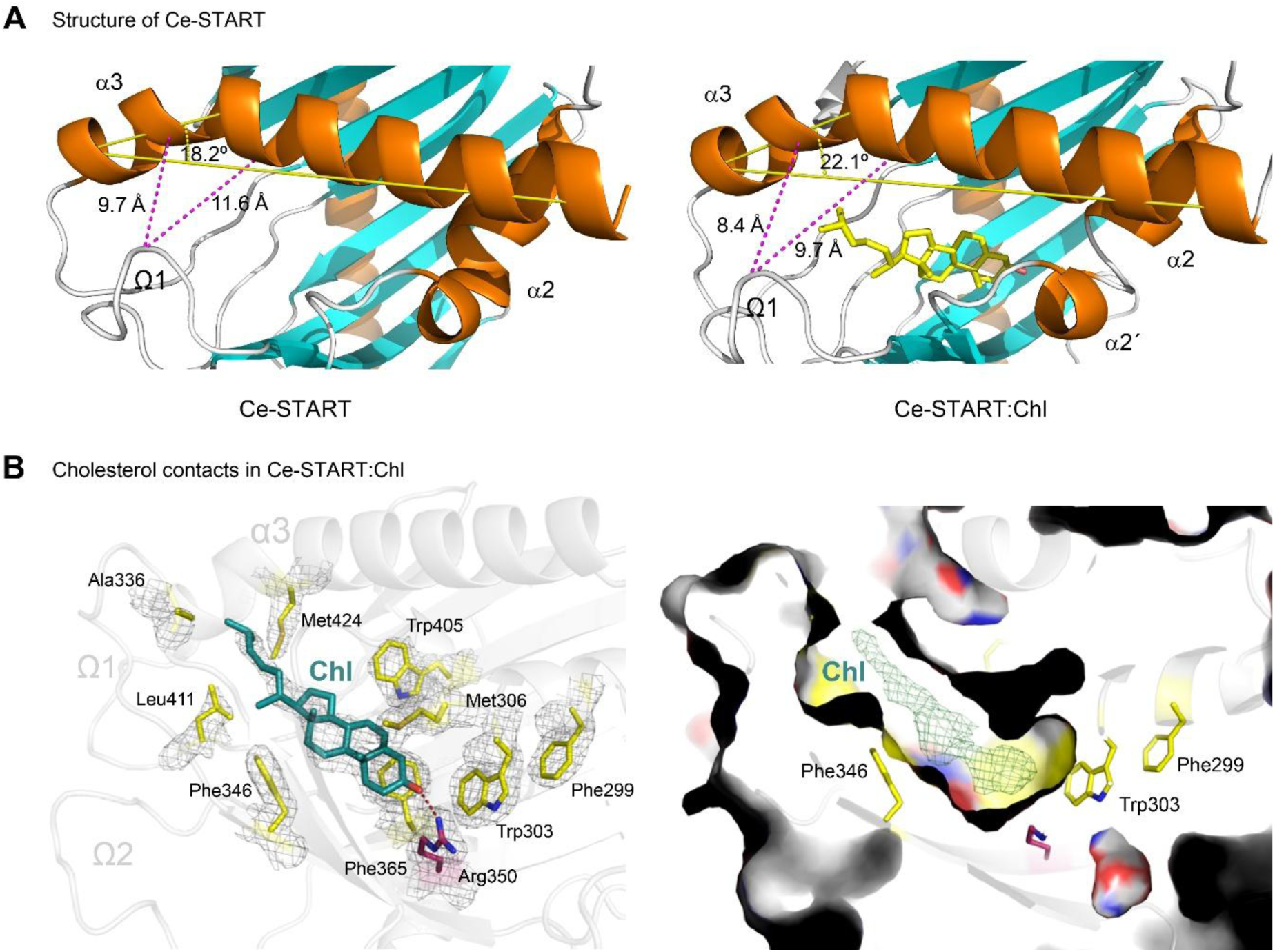
Cholesterol binding to Ce-START elicits conformational changes. **(A)** Comparison of the hydrophobic cavities of Ce-START and Ce-START:Chl complex. The distances between alanine 336 (Ω1) and glycine 423 and tyrosine 427 (both in the α3 helix) are indicated by magenta dashed lines. The torsion angle of the α3 helix is shown in yellow. **(B)** Close-up view of the cholesterol binding site in Ce-START (chain C), shown in two complementary representations. On the left, hydrophobic amino acid residues located within a 4 Å radius of the cholesterol molecule (green) are displayed as sticks and highlighted in yellow; these residues establish non-polar interactions with the hydrophobic portion of the cholesterol molecule, promoting its stabilization within the cavity. Arginine residue 350, shown in red, forms a polar interaction with the hydroxyl group of cholesterol, indicated by a red dashed line. The *2mFo-DFc* electron density map (gray mesh), contoured at 1 σ, is superimposed onto the highlighted residues and the cholesterol molecule. On the right, the same binding site is displayed with the protein rendered as a molecular surface; the cholesterol molecule was omitted from the model and the corresponding omit *mFo-DFc* difference electron density (green mesh), contoured at 2.75 σ, is shown within the cavity, confirming the presence and placement of the ligand.

Previous analyses of human START domains that interact with cholesterol identified a set of amino acid residues that are critical for proper ligand binding (19,20,24). In the Ce-START structure, we found that some of these residues, such as Met306 and Arg350, are conserved (**Fig. 3B** and **Supplementary Fig. 1**). In particular, Arg350 appears to play a crucial role in sterol binding, as it would stabilize the charge of the cholesterol hydroxyl group through a hydrogen bond with its side chain, located 3.1 Å from the substituent group (**Fig. 3B**). Finally, our results show that stabilization of cholesterol at the binding site is also mediated by contacts between the lipid molecule and residues of Ce-START, such as Met306, Val313, Phe346, Phe365, Trp405, Leu411 and Met424. All of them belong to the same sub-family of non-polar amino acids, stressing their role in stabilizing the hydrophobic half of the cholesterol molecule at the binding site.

### Physiological role of STARD3

*C. elegans* is an excellent model for studying cholesterol trafficking and metabolism because this nematode is unable to synthesize cholesterol and requires its presence in the diet. This allows an easy manipulation of worm sterol pools by introducing variations of its levels in the diet (34). To study the physiological role of STARD3 in cholesterol transport, we generated a *stard3-*(*-*) mutant by deleting 2204 bp from the *tag-340* gene (**Supplementary Fig. 6**). We then cultured the worms on cholesterol-free plates and assessed their development up to the adult stage. In the first generation (F1) of animals grown without cholesterol, no differences were observed between *stard3* mutants and the N2 wild-type strain, as most individuals reached adulthood (61% and 74% for wild type N2 and *stard3-*(*-*), respectively, *p* > 0.05, Kruskal-Wallis test followed for Dunn’s multiple comparisons test, **Fig. 4A**). However, when worms were cultured for two consecutive generations (F2) in the absence of cholesterol, virtually all *stard3-*(*-*) mutants failed to complete reproductive development and arrested at a stage resembling L2 (L2*). In contrast, a statistically significant proportion of N2 worms developed to adulthood under the same conditions (10%, *p* < 0.05, Kruskal-Wallis test followed by Dunn’s multiple comparisons test, **Fig. 4B, C**). The arrest phenotype was partially reversed by supplementing the medium with cholesterol, resulting in recovery in both strains, with no significant differences between them (*p* > 0.05, Kruskal-Wallis test followed by Dunn’s multiple comparisons test, **Fig. 4B**).

**Figure 4.**
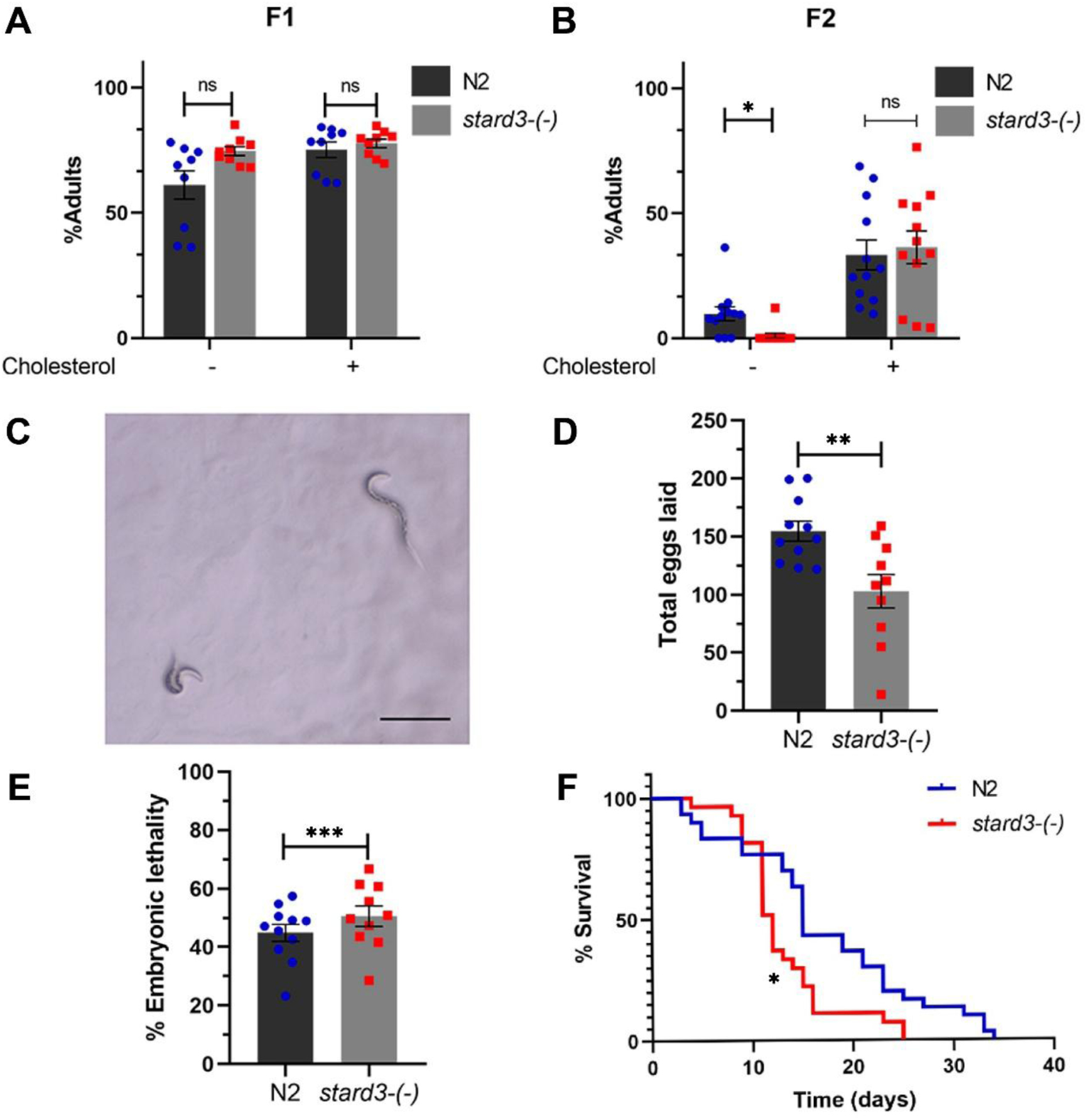
STARD3 becomes physiologically relevant when cholesterol availability is limited. **(A, B)** Average percentage of adult worms reached by N2 and *stard3-*(*-*) animals grown in the presence or absence of cholesterol at 20°C. **(A)** worms grown for one generation (F1) without cholesterol and observed 72 hours after embryo hatching. **(B)** Worms grown for two consecutive generations (F2) without cholesterol and observed 72 hours after embryo hatching. Bars represent means, and error bars represent SEM. n = 3 independent experiments. **(C)** Representative image of *stard3-*(*-*) L2 worms grown for two generations on cholesterol-free medium, observed 72 hours after embryo hatching. Scale bar, 200 µm. **(D)** Mean number of eggs laid (brood size), and **(E)** percentage of embryonic lethality for *stard3-*(*-*) and N2 worms after growing for one generation in the absence of cholesterol at 20°C. Error bars represent SEM. A minimum of 10 F1 worms were analyzed for each condition. **(F)** Kaplan–Meier survival curve of N2 and *stard3-*(*-*) worms grown on standard NGM medium without cholesterol deprivation. Viability was assessed every 24 hours and expressed as the percentage of surviving worms.

To evaluate the role of *stard3-*(*-*) in fertility under low dietary cholesterol, we assessed brood size and embryonic lethality in the first generation of mutant worms grown without exogenous sterols. *stard3-*(*-*) mutants exhibited a 34% reduction in brood size compared with N2 worms, accompanied by a statistically significant increase in embryonic lethality (*p* < 0.01 and *p* < 0.001, respectively, Welch’s unpaired t-test and Fisher’s exact test, **Fig. 4D-E**). Under standard culture conditions, fertility was not affected (*p* > 0.05, unpaired t-test, **Supplementary Fig. 7**). However, in this same medium, *stard3-*(*-*) mutants exhibited reduced survival compared to the wild-type strain, with median lifespans of 12 and 15 days, respectively, (*p* < 0.05, Log-Rank/Mantel Cox test, **Fig. 4F**). These results suggest that STARD3 is not essential for cholesterol transport but may act synergistically with other sterol mobilization systems in the worm. The failure to thrive in the second generation without exogenous cholesterol, together with the decreased brood size observed in the first generation under cholesterol deprivation, indicates that STARD3’s transporter function becomes increasingly critical when environmental cholesterol levels decrease.

Several proteins, including the NPC1-NPC2 complex, ORP5, and STARD3, participate in cholesterol transport from late endosomes/lysosomes (LE/Ly) (17,18,47). Although previous studies in cell culture have shown that STARD3 facilitates the redistribution of intracellular cholesterol from the ER to LE/Ly, it cannot be ruled out that this mechanism may also function in the reverse direction under specific regulatory conditions (17,47). Moreover, there is evidence suggesting that STARD3, or another lipid transfer protein, may mediate sterol transport even in the absence of functional NPC1 (48). To assess the role of STARD3 in sterol trafficking through the NPC system, we silenced *stard3* by RNAi in strain JT10800, a double mutant for *ncr-2*(*-*); *ncr-1*(*-*), the *C. elegans* orthologs of NPC1 (49,50). This strain forms dauers due to its inability to export cholesterol from LE/Ly (6,11,33). We hypothesized that if STARD3 functions similarly, or in parallel to the NPC system by promoting cholesterol efflux from LE/Ly, its absence should exacerbate the dauer phenotype. Indeed, silencing *stard3* increased the number of worms entering the dauer stage compared to controls (56% and 76% for EV and *stard3*RNAi, respectively, *p* < 0.01, Mann-Whitney U test, **Fig. 5A**). Since dauer formation results from the inability of worms to synthesize dafachronic acids (DAs) from cholesterol reservoirs, we hypothesized that STARD3 depletion could exacerbate cholesterol deficiency by limiting its availability at DA production sites. To test this hypothesis, we evaluated whether higher cholesterol availability in the growth medium could compensate for STARD3 silencing in the *ncr-2*(*-*)*; ncr-1*(*-*) mutant. We observed that, although a ten-fold increase in cholesterol concentration reduced dauer formation in *ncr-2-*(*-*)*; ncr-1-*(*-*) mutants, this excess was unable to rescue the phenotype when *stard3* was silenced (*p* < 0.05 and *p* > 0.05, respectively, Kruskal-Wallis test followed by Dunn’s multiple comparisons test, **Fig. 5B**). Taken together, these results suggest that STARD3 in *C. elegans* functions analogously to the NPC system to enable cholesterol efflux from LE/Ly, and is important for proper delivery to sites involved in DA biosynthesis.

**Figure 5.**
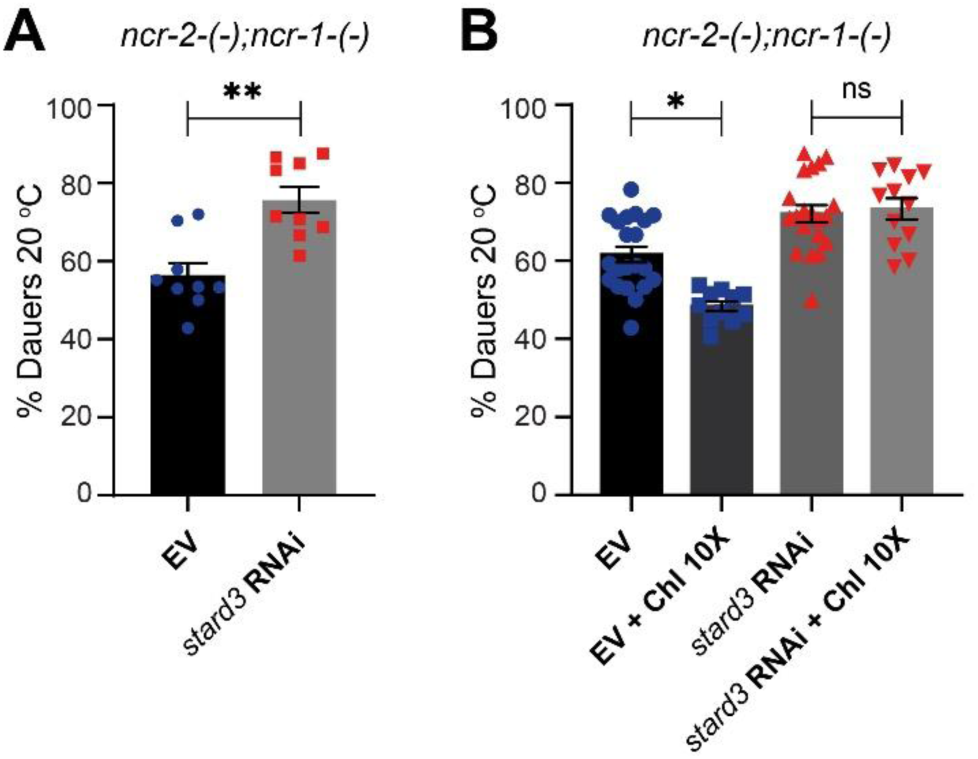
*C. elegans* STARD3 acts analogously to NCR-2 and NCR-1. Average percentage of dauer larvae in JT10800 worms [*ncr-2*(*2022*)*; ncr-1*(*2023*)] maintained at 20 °C and fed with *E. coli* HT115 transformed with an empty vector (EV) or with a vector expressing RNAi against *stard3* (*stard3* RNAi). Experiments were done with standard cholesterol supplementation (13 µM) **(A)** and repeated in the presence of 10X cholesterol (130 µM) **(B)**. Data from at least three independent experiments.

## Discussion

In this work, we characterized the conformational properties of the START domain of *C. elegans* STARD3 using biophysical approaches. We determined that this domain is folded in its native state and has the ability to bind cholesterol with high affinity and a 1:1 stoichiometry, similar to its human ortholog. Additionally, we solved the structure of Ce-START by X-ray protein crystallography, both in its free form and bound to cholesterol. By comparing these structures with human START domain structures available from previous works, we confirmed that the START domain of *C. elegans* is highly similar to its human orthologs (13,19,20,44–46). This finding is consistent with the high conservation of START domains among eukaryotic species (13,19).

The structure of Ce-START in complex with cholesterol provided empirical evidence about the importance of key residues, such as Arg350, in ligand binding. According to the proposed model for the START domain of human STARD3 based on molecular docking studies, conformational changes in the Ω-loop (loop Ω 1) would induce displacement of the α3 helix, thereby facilitating the opening of the hydrophobic cavity and allowing cholesterol entry or release (20,25). In our structure of cholesterol-bound Ce-START, we observed that the Ω-loop moves closer to the α3 helix compared to the apo state of Ce-START. We also detected an increase in the torsion angle of this helix after ligand binding. The structural transitions elicited by cholesterol binding to Ce-START are consistent with the proposed model and provide direct empirical support for the underlying mechanism. The structural differences observed between the free and bound states of Ce-START allow us to hypothesize that local conformational changes occur in the structures forming the roof of the hydrophobic cavity, enabling cholesterol binding and stabilization of the complex. Furthermore, the hydrophobic residues of the Ω-loop could serve as anchoring points for the START domain to membranes from which it captures or releases cholesterol. Thus, the observed structural changes would be key to enable efficient cholesterol transport, although further experiments are required to confirm this hypothesis.

We also observed that methionine residues at positions 306 and 424 lie within the hydrophobic cavity and in close proximity to the cholesterol molecule. Previous work has shown that the START domain of human STARD3 participates in the detoxification of cholesterol hydroperoxides at the endosomal membrane. This activity relies on the direct binding of hydroperoxides to human START and the involvement of Met307 and Met427 in oxidation–reduction cycles mediated by the methionine sulfoxide reductase protein family (51,52). The structural homology between the human and *C. elegans* START domains, together with the conservation Met306 and Met424 and their proximity to the cholesterol molecule, supports those findings and suggests that a similar mechanism may operate in *C. elegans*, which also possesses active methionine sulfoxide reductase proteins (53). Such a mechanism could be particularly relevant for *C. elegans* due to its cholesterol-auxotrophic nature and the strict requirement to maintain appropriate cholesterol levels for normal development, growth, and reproduction.

Phenotypic analysis of the *stard3-*(*-*) mutant under cholesterol-restricted conditions revealed no differences from the N2 wild type during the first generation. However, brood size and embryonic viability were significantly reduced in *stard3-*(*-*) animals, consistent with previous reports showing that cholesterol deficiency causes multiple germline defects in *C. elegans* (54,55). In the second generation, cholesterol-deprived mutants exhibited complete larval arrest, whereas control worms showed only a small proportion of arrested larvae. The rescue of this phenotype by exogenous cholesterol supplementation indicates that STARD3 is essential for efficient sterol mobilization and homeostasis when internal cholesterol reserves are depleted.

Our results show that STARD3 deficiency in *ncr-2-*(*-*)*;ncr-1-*(*-*) background increases dauer formation by 20% at 20°C, suggesting that this transporter plays a key role in regulating cholesterol trafficking in *C. elegans*. It is known that the *ncr-2-*(*-*)*;ncr-1-*(*-*) strain lacks the NCR-1 transporter, homologous to human NPC1(11,33), which facilitates intracellular cholesterol trafficking (49). The absence of NCR-1 leads to cholesterol accumulation in LE/Ly compartments, making it unavailable for DA synthesis and inducing dauer arrest (49).

STARD3 has been identified as a cholesterol transporter in endosomal compartments in other organisms, although its role in *C. elegans* had not been previously studied. Our findings suggest that STARD3 functions independently of the NCR system and that its activity is required for normal longevity and for attenuating the dauer phenotype in NCR-1–deficient worms. This allows us to hypothesize that STARD3 facilitates cholesterol mobilization from LE/Ly to other cellular compartments, ensuring its availability for key processes involved in dauer exit (6,56), specially under conditions of cholesterol deprivation. The redundancy of these pathways highlights the importance of cholesterol transport in animals and explains why *stard3-*(*-*) worms do not exhibit a dauer phenotype under normal growth conditions.

The function of STARD3 as a transporter could represent a relevant mechanism in the regulation of sterol metabolism in *C. elegans* and in coordinating the cellular events necessary for initiating reproductive development. Although additional experiments are needed to confirm these hypotheses, our structural and biological data suggest that STARD3-mediated, non-vesicular cholesterol transport is conserved from *C. elegans* to higher organisms, including humans. This finding further supports the value of this animal model for elucidating the role of cholesterol metabolism in both physiological and pathological processes.

## Data availability

All relevant data are included in the article and its supporting information. PDB files for Ce-START and Ce-START:Chl structures can be accessed from the Protein Data Bank using the codes 13FT, and 13FR, respectively.

## Supplemental data

This article contains supplemental data

## Acknowledgements

This research was supported by the Richard Lounsbery Foundation (RLF-2021 to A. B. and D. d. M), Agencia I+D+i (PICT-2018-02572 to A.B. and D. d. M.), ASaCTeI (PEICID-2022-232 to M.C.M.) and CONICET. B.B., A.S. and F.A.B. received fellowships from CONICET. V.A.L., M.C.M., D.A., M.N.L., D.d.M. and A.B. are staff members of CONICET. We thank Alejandro Buschiazzo, Nicole Larrieux, and Albertina Scattolini for assistance with crystal handling and diffraction data collection. We also thank the Diamond Light Source for access to the X-ray beamline facilities. We thank Roberta Pierattelli for helpful discussions. Further support has been provided from access to the CERM/CIRMMP NMR research Infrastructure within the EC Horizon 2020 iNEXT-Discovery project (PID 18340). We acknowledge Andrea Coscia and Alejandro Gago for maintenance of the NMR infrastructure and Cecilia Vranych for assistance with *C. elegans* growth and maintenance.

## Author Contributions

B.B., D.d.M. and A.B. crafted the main hypothesis and designed research. D.d.M. and A.B. were responsible for project funding and administration. B.B., A.S., F.A.B., V.A.L., M.C.M., D.d.M and A.B. performed molecular biology, biochemistry and *C. elegans* experiments. B.B., F.A.B. and A.B. performed CD, fluorescence and NMR experiments. B.B., F.A.B. and A.B. modeled proteins structures. B.B. D.A. and M.N.L. crystalized the proteins and solved the X-Ray structures. B.B., D.d.M. and A.B. wrote the paper, and all authors discussed the results and commented on the manuscript.

## Funding sources

This work was funded by CONICET, Agencia I+D+i (PICT-2018-02572 to A.B. and D. d. M.) and the Richard Lounsbery Foundation (2021 to A. B. and D. d. M).

## Abbreviations

*C. elegans*: *Caenorhabditis elegans*
StAR: steroidogenic acute regulatory proteins
STARD3: StAR-related lipid transfer protein 3
Ce-START: START domain of *C. elegans* STARD3
DA: dafachronic acids
NPC1: Niemann-Pick type C1
LE/Ly: late endosomes/lysosomes
ER: endoplasmic reticulum
VAP: VAMP-associated proteins
FFAT: two phenylalanines in an acidic tract
ChlCDX: water-soluble cholesterol
mCDX: methyl-β-cyclodextrin
NMR: nuclear magnetic resonance
HSQC: heteronuclear single quantum coherence
CD: circular dichroism
NGM: nematode growth medium
*Ka*: affinity constant
*Kd*: dissociation constant
RMSD: Root Mean Square Deviation
NMR: nuclear magnetic resonance
HSQC: heteronuclear single quantum coherence

## Notes

### Competing Interest Statement

The authors have declared no competing interest.

### Summary of Updates

New X-ray analysis of the Ce-START:cholesterol complex

## References

1. Ikonen, E. (2008) Cellular cholesterol trafficking and compartmentalization. Nat Rev Mol Cell Biol 9, 125–138

2. Maxfield, F. R., and van Meer, G. (2010) Cholesterol, the central lipid of mammalian cells. Curr Opin Cell Biol 22, 422–429

3. Brenner, S. (1974) The genetics of *Caenorhabditis elegans*. Genetics 77, 71–94

4. Watts, J. L., and Ristow, M. (2017) Lipid and Carbohydrate Metabolism in *Caenorhabditis elegans*. Genetics 207, 413–446

5. Rauthan, M., and Pilon, M. (2011) The mevalonate pathway in *C. elegans*. Lipids Health Dis 10, 243

6. Fielenbach, N., and Antebi, A. (2008) *C. elegans* dauer formation and the molecular basis of plasticity. Genes Dev 22, 2149–2165

7. Gerisch, B., and Antebi, A. (2004) Hormonal signals produced by DAF-9/cytochrome P450 regulate *C. elegans* dauer diapause in response to environmental cues. Development 131, 1765–1776

8. Jeong, M. H., Kawasaki, I., and Shim, Y. H. (2010) A circulatory transcriptional regulation among daf-9, daf-12, and daf-16 mediates larval development upon cholesterol starvation in *Caenorhabditis elegans*. Dev Dyn 239, 1931–1940

9. Kimble, J., and Sharrock, W. J. (1983) Tissue-specific synthesis of yolk proteins in *Caenorhabditis elegans*. Dev Biol 96, 189–196

10. Lu, A. (2022) Endolysosomal cholesterol export: More than just NPC1. Bioessays 44, e2200111

11. Li, J., Brown, G., Ailion, M., Lee, S., and Thomas, J. H. (2004) NCR-1 and NCR-2, the *C. elegans* homologs of the human Niemann-Pick type C1 disease protein, function upstream of DAF-9 in the dauer formation pathways. Development 131, 5741–5752

12. Zhang, M. G., and Sternberg, P. W. (2022) Both entry to and exit from diapause arrest in *Caenorhabditis elegans* are regulated by a steroid hormone pathway. Development 149, dev200173

13. Clark, B. J. (2020) The START-domain proteins in intracellular lipid transport and beyond. Mol Cell Endocrinol 504, 110704

14. Alpy, F., Stoeckel, M. E., Dierich, A., Escola, J. M., Wendling, C., Chenard, M. P., et al. (2001) The steroidogenic acute regulatory protein homolog MLN64, a late endosomal cholesterol-binding protein. J Biol Chem 276, 4261–4269

15. Torres, S., Balboa, E., Zanlungo, S., Enrich, C., Garcia-Ruiz, C., and Fernandez-Checa, J. C. (2017) Lysosomal and Mitochondrial Liaisons in Niemann-Pick Disease. Front Physiol 8, 982

16. Voilquin, L., Lodi, M., Di Mattia, T., Chenard, M.-P., Mathelin, C., Alpy, F., et al. (2019) STARD3: A Swiss Army Knife for Intracellular Cholesterol Transport. Contact 2, 1–15

17. Wilhelm, L. P., Wendling, C., Vedie, B., Kobayashi, T., Chenard, M. P., Tomasetto, C., et al. (2017) STARD3 mediates endoplasmic reticulum-to-endosome cholesterol transport at membrane contact sites. EMBO J 36, 1412–1433

18. Luo, J., Jiang, L. Y., Yang, H., and Song, B. L. (2019) Intracellular Cholesterol Transport by Sterol Transfer Proteins at Membrane Contact Sites. Trends Biochem Sci 44, 273–292

19. Thorsell, A. G., Lee, W. H., Persson, C., Siponen, M. I., Nilsson, M., Busam, R. D., et al. (2011) Comparative structural analysis of lipid binding START domains. PLoS One 6, e19521

20. Tsujishita, Y., and Hurley, J. H. (2000) Structure and lipid transport mechanism of a StAR-related domain. Nat Struct Biol 7, 408–414

21. Watari, H., Arakane, F., Moog-Lutz, C., Kallen, C. B., Tomasetto, C., Gerton, G. L., et al. (1997) MLN64 contains a domain with homology to the steroidogenic acute regulatory protein (StAR) that stimulates steroidogenesis. Proc Natl Acad Sci U S A 94, 8462–8467

22. Robert, X., and Gouet, P. (2014) Deciphering key features in protein structures with the new ENDscript server. Nucleic Acids Res 42, W320–324

23. Mirdita, M., Schutze, K., Moriwaki, Y., Heo, L., Ovchinnikov, S., and Steinegger, M. (2022) ColabFold: making protein folding accessible to all. Nat Methods 19, 679–682

24. Letourneau, D., Lorin, A., Lefebvre, A., Cabana, J., Lavigne, P., and LeHoux, J. G. (2013) Thermodynamic and solution state NMR characterization of the binding of secondary and conjugated bile acids to STARD5. Biochim Biophys Acta 1831, 1589–1599

25. Roostaee, A., Barbar, E., Lehoux, J. G., and Lavigne, P. (2008) Cholesterol binding is a prerequisite for the activity of the steroidogenic acute regulatory protein (StAR). Biochem J 412, 553–562

26. Kabsch, W. (2010) Xds. Acta Crystallogr D Biol Crystallogr 66, 125–132

27. Evans, P. R., and Murshudov, G. N. (2013) How good are my data and what is the resolution? Acta Crystallogr D Biol Crystallogr 69, 1204–1214

28. Winn, M. D., Ballard, C. C., Cowtan, K. D., Dodson, E. J., Emsley, P., Evans, P. R., et al. (2011) Overview of the CCP4 suite and current developments. Acta Crystallogr D Biol Crystallogr 67, 235–242

29. McCoy, A. J., Grosse-Kunstleve, R. W., Adams, P. D., Winn, M. D., Storoni, L. C., and Read, R. J. (2007) Phaser crystallographic software. J Appl Crystallogr 40, 658–674

30. Emsley, P., Lohkamp, B., Scott, W. G., and Cowtan, K. (2010) Features and development of Coot. Acta Crystallogr D Biol Crystallogr 66, 486–501

31. Afonine, P. V., Grosse-Kunstleve, R. W., Echols, N., Headd, J. J., Moriarty, N. W., Mustyakimov, M., et al. (2012) Towards automated crystallographic structure refinement with phenix.refine. Acta Crystallogr D Biol Crystallogr 68, 352–367

32. Williams, C. J., Headd, J. J., Moriarty, N. W., Prisant, M. G., Videau, L. L., Deis, L. N., et al. (2018) MolProbity: More and better reference data for improved all-atom structure validation. Protein Sci 27, 293–315

33. Galles, C., Prez, G. M., Penkov, S., Boland, S., Porta, E. O. J., Altabe, S. G., et al. (2018) Endocannabinoids in *Caenorhabditis elegans* are essential for the mobilization of cholesterol from internal reserves. Sci Rep 8, 6398

34. Matyash, V., Entchev, E. V., Mende, F., Wilsch-Brauninger, M., Thiele, C., Schmidt, A. W., et al. (2004) Sterol-derived hormone(s) controls entry into diapause in *Caenorhabditis elegans* by consecutive activation of DAF-12 and DAF-16. PLoS Biol 2, e280

35. Ashrafi, K., Chang, F. Y., Watts, J. L., Fraser, A. G., Kamath, R. S., Ahringer, J., et al. (2003) Genome-wide RNAi analysis of *Caenorhabditis elegans* fat regulatory genes. Nature 421, 268–272

36. Fraser, A. G., Kamath, R. S., Zipperlen, P., Martinez-Campos, M., Sohrmann, M., and Ahringer, J. (2000) Functional genomic analysis of *C. elegans* chromosome I by systematic RNA interference. Nature 408, 325–330

37. Conte, D., Jr., MacNeil, L. T., Walhout, A. J. M., and Mello, C. C. (2015) RNA Interference in *Caenorhabditis elegans*. Curr Protoc Mol Biol 109, 26 23 21–26 23 30

38. Miotto, M. C., Binolfi, A., Zweckstetter, M., Griesinger, C., and Fernandez, C. O. (2014) Bioinorganic chemistry of synucleinopathies: deciphering the binding features of Met motifs and His-50 in AS-Cu(I) interactions. J Inorg Biochem 141, 208–211

39. Becker, W., Bhattiprolu, K. C., Gubensak, N., and Zangger, K. (2018) Investigating Protein-Ligand Interactions by Solution Nuclear Magnetic Resonance Spectroscopy. Chemphyschem 19, 895–906

40. Frijlink, H. W., Eissens, A. C., Hefting, N. R., Poelstra, K., Lerk, C. F., and Meijer, D. K. (1991) The effect of parenterally administered cyclodextrins on cholesterol levels in the rat. Pharm Res 8, 9–16

41. Lakowicz, J. R. (ed) (2006) Protein Fluorescence, Springer, Boston, MA.

42. Jarmoskaite, I., AlSadhan, I., Vaidyanathan, P. P., and Herschlag, D. (2020) How to measure and evaluate binding affinities. Elife 9, e57264

43. Greenfield, N. J. (2006) Using circular dichroism spectra to estimate protein secondary structure. Nat Protoc 1, 2876–2890

44. Horvath, M. P., George, E. W., Tran, Q. T., Baumgardner, K., Zharov, G., Lee, S., et al. (2016) Structure of the lutein-binding domain of human StARD3 at 1.74 A resolution and model of a complex with lutein. Acta Crystallogr F Struct Biol Commun 72, 609–618

45. Romanowski, M. J., Soccio, R. E., Breslow, J. L., and Burley, S. K. (2002) Crystal structure of the Mus musculus cholesterol-regulated START protein 4 (StarD4) containing a StAR-related lipid transfer domain. Proc Natl Acad Sci U S A 99, 6949–6954

46. Tan, L., Tong, J., Chun, C., and Im, Y. J. (2019) Structural analysis of human sterol transfer protein STARD4. Biochem Biophys Res Commun 520, 466–472

47. Hoglinger, D., Burgoyne, T., Sanchez-Heras, E., Hartwig, P., Colaco, A., Newton, J., et al. (2019) NPC1 regulates ER contacts with endocytic organelles to mediate cholesterol egress. Nat Commun 10, 4276

48. Martello, A., Platt, F. M., and Eden, E. R. (2020) Staying in touch with the endocytic network: The importance of contacts for cholesterol transport. Traffic 21, 354–363

49. Rosenbaum, A. I., and Maxfield, F. R. (2011) Niemann-Pick type C disease: molecular mechanisms and potential therapeutic approaches. J Neurochem 116, 789–795

50. Sym, M., Basson, M., and Johnson, C. (2000) A model for niemann-pick type C disease in the nematode *Caenorhabditis elegans*. Curr Biol 10, 527–530

51. Lim, J. M., Lim, J. C., Kim, G., and Levine, R. L. (2018) Myristoylated methionine sulfoxide reductase A is a late endosomal protein. J Biol Chem 293, 7355–7366

52. Lim, J. M., Sabbasani, V. R., Swenson, R. E., and Levine, R. L. (2023) Methionine sulfoxide reductases and cholesterol transporter STARD3 constitute an efficient system for detoxification of cholesterol hydroperoxides. J Biol Chem 299, 105099

53. Minniti, A. N., Cataldo, R., Trigo, C., Vasquez, L., Mujica, P., Leighton, F., et al. (2009) Methionine sulfoxide reductase A expression is regulated by the DAF-16/FOXO pathway in *Caenorhabditis elegans*. Aging Cell 8, 690–705

54. Merris, M., Wadsworth, W. G., Khamrai, U., Bittman, R., Chitwood, D. J., and Lenard, J. (2003) Sterol effects and sites of sterol accumulation in *Caenorhabditis elegans*: developmental requirement for 4alpha-methyl sterols. J Lipid Res 44, 172–181

55. Shim, Y. H., Chun, J. H., Lee, E. Y., and Paik, Y. K. (2002) Role of cholesterol in germ-line development of *Caenorhabditis elegans*. Mol Reprod Dev 61, 358–366

56. Antebi, A., Culotti, J. G., and Hedgecock, E. M. (1998) daf-12 regulates developmental age and the dauer alternative in *Caenorhabditis elegans*. Development 125, 1191–1205

