## Supporting Information for "STARD3 mediates non-vesicular cholesterol transport in *Caenorhabditis elegans*"

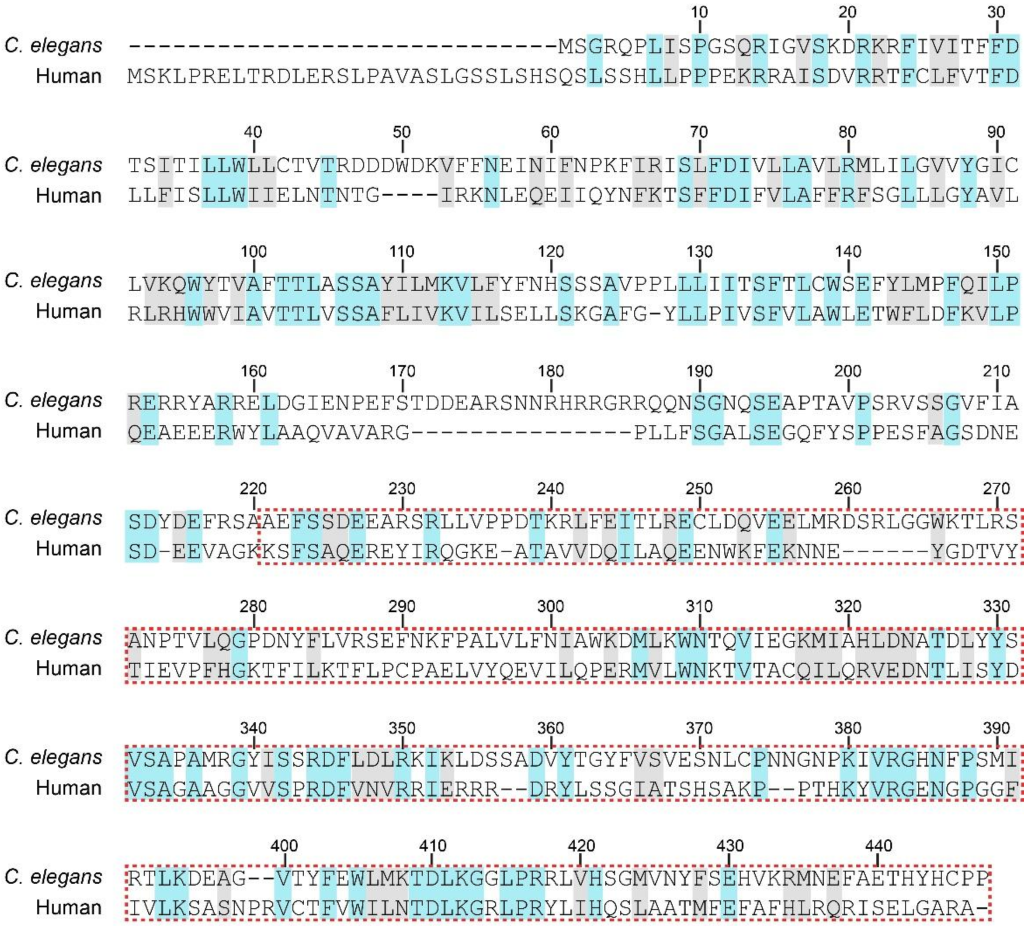


**Supplementary figure 1.** Primary sequence alignments between *C. elegans* STARD3 (top, UniProt Q19819) and human STARD3 (bottom, UniProt Q14849). Identical and similar residues are shaded in cyan and gray, respectively. Red dashed boxes indicate the amino acid sequence of *C. elegans* START domain used in this work. The purified sequence contains an extra G residue at the N-terminus from the TEV cleavage site.

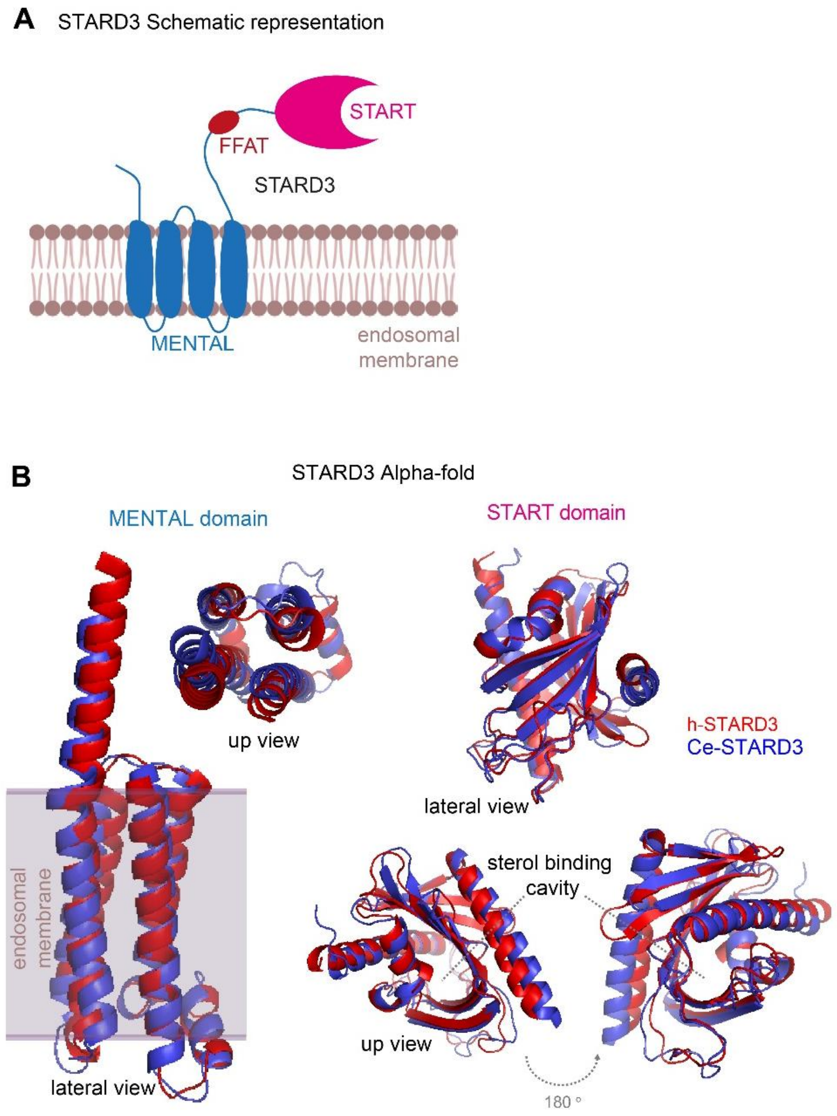


**Supplementary Figure 2.** **(A)** Schematic representation of STARD3 protein. The START, FFAT and MENTAL domains are depicted in pink, red and cyan, respectively. **(B)** Overlay of the AlphaFold-predicted 3D structures of human STARD3 (red) and Ce-STARD3 (blue). MENTAL domain is illustrated within the endosomal membrane.

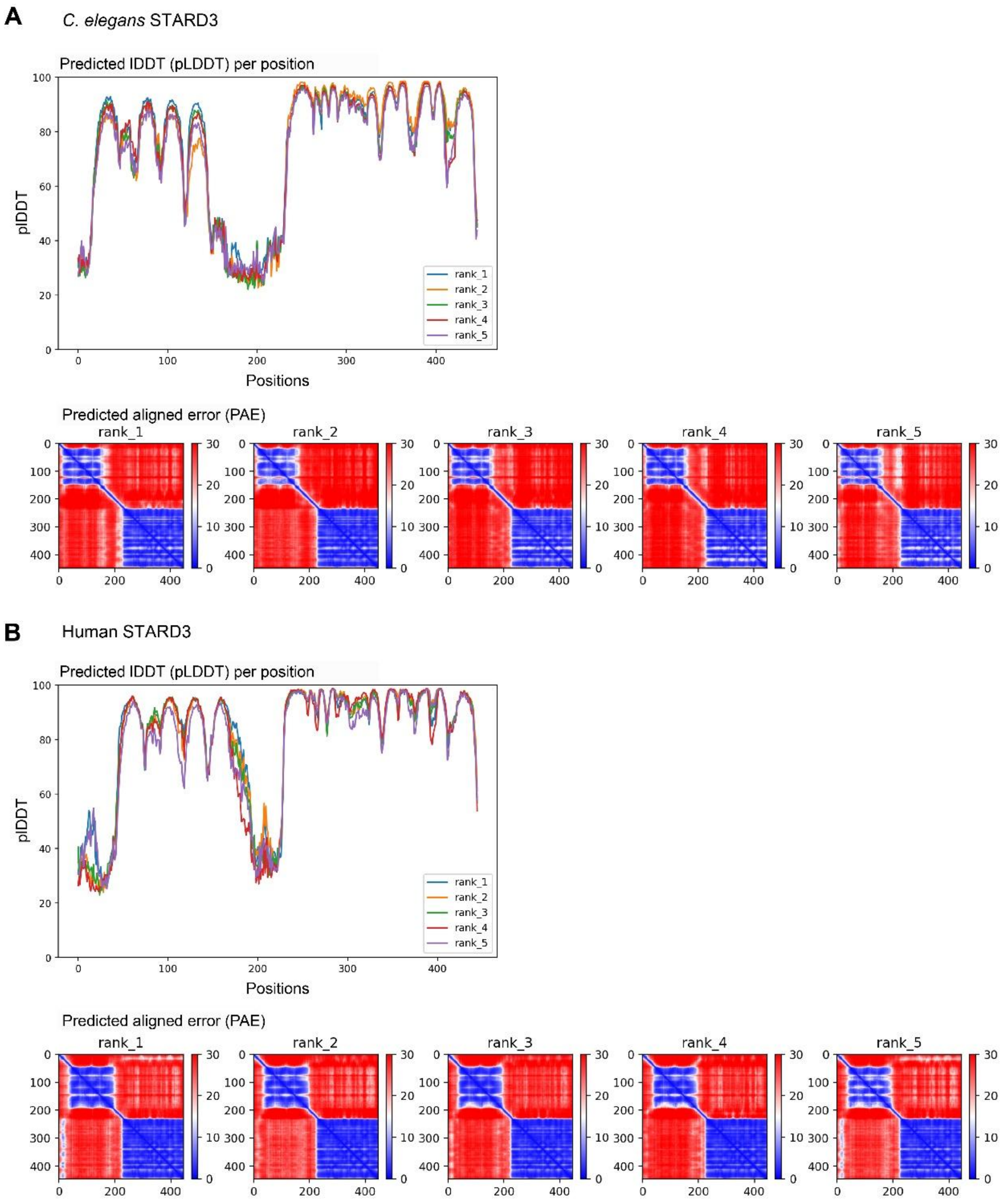


**Supplementary Figure 3.** Predicted local distance difference test (pLDDT) scores (top) and predicted aligned error (PAE) (bottom) for the AlphaFold2 models of *C. elegans* (**A**) and human (**B**) STARD3 proteins shown in **Supplementary Fig. 2b**.

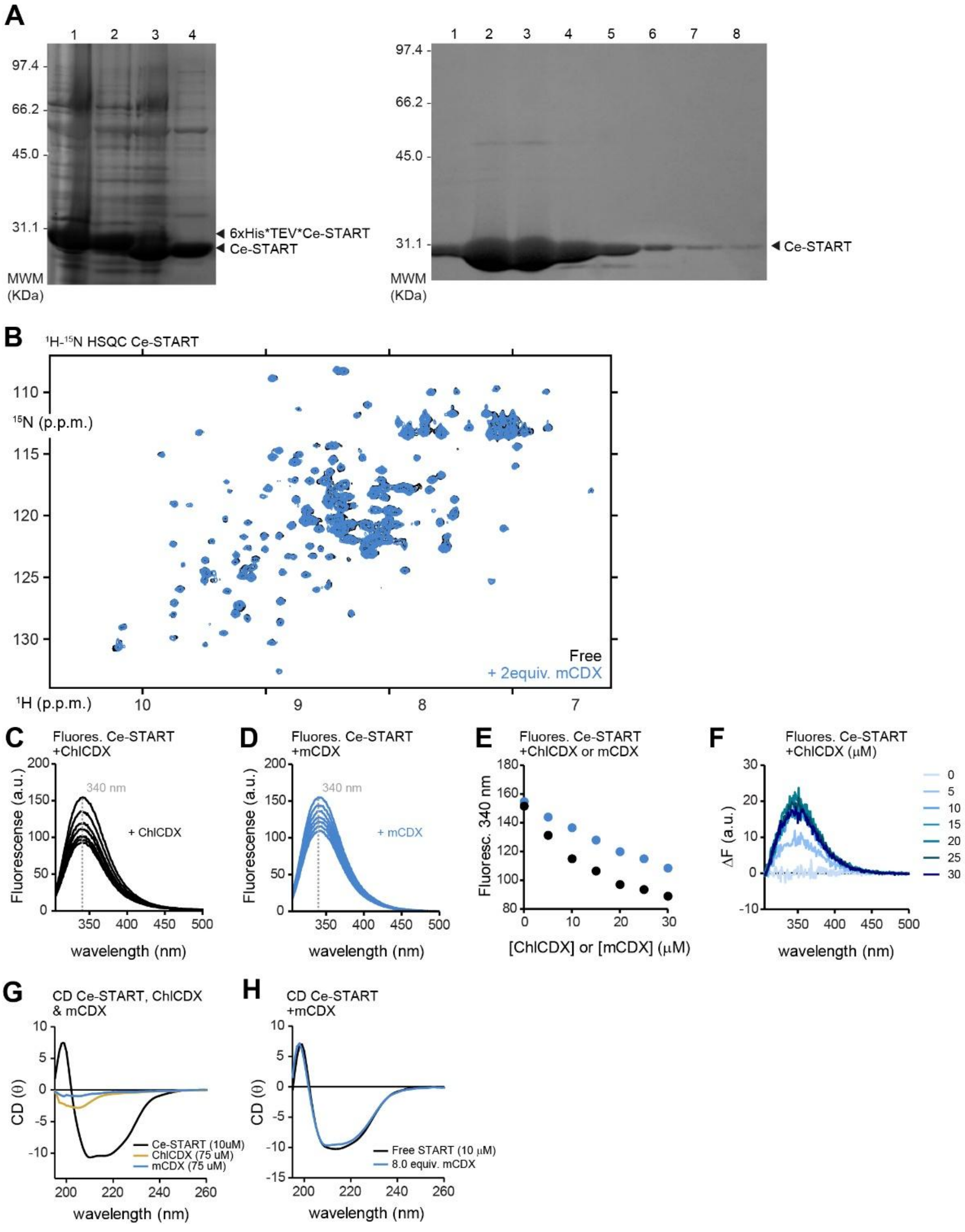


**Supplementary Figure 4. (A)** SDS-PAGE analysis of the purification of recombinant Ce-START. Left panel shows the removal of the 6xHis-TEV tag. Lane 1, corresponds to 6xHis-TEV-Ce-START; lane 2 6xHis-TEV-Ce-START + TEV protease before cleavage; lane 3 6xHis-TEV-Ce-START + TEV protease after cleavage and lane 4 Ce-START purified from the 6xHis-TEV tag using Ni-NTA resin. Right panel shows, the SEC elution profile of Ce-START. Fractions, 2-5 were pooled and concentrated for further analysis. The mature protein contains a Gly at position 1 from the TEV processing site. **(B)** Overlay of $^{1}H$-$^{15}N$ HSQC spectra of 150 μM, $^{15}N$-isotopically enriched Ce-START in the absence (black) and presence (cyan) of 2 equivalents of cholesterol-free methyl-cyclodextrin (mCDX). This corresponds to the same amount of mCDX complex to cholesterol **(C, D)** Intrinsic Trp fluorescence spectra of 10 μM Ce-START in presence of cholesterol-mCDX **(C)** or mCDX **(D)**. **(E)** Intrinsic Trp fluorescence of 10 μM Ce-START at increasing concentrations of ChlDX and mCDX. Fluorescence intensity maxima at 340 nm were obtained from the experiments showed in panels **(C, D). (F)** Difference fluorescence spectra of Ce-START-Chl complex after subtraction of the mCDX contribution from each added concentration. **(G)** Far-UV CD spectra of 8 μM Ce-START and of 75 μM of Chl:CDX (yellow) and 75 mM μCDX (cyan). **(H)** Far-UV CD spectra of 8 μM Ce-START in the absence (black line) or presence of 8 equivalents of mCDX (cyan).

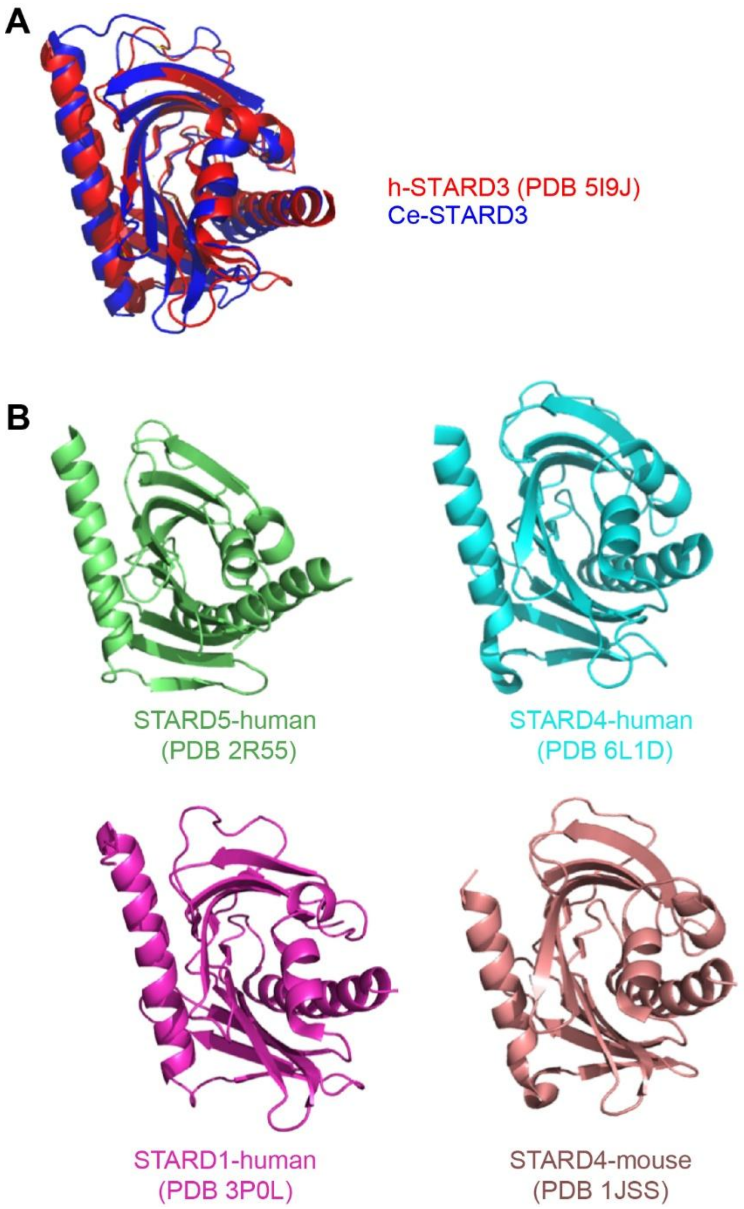


**Supplementary Figure 5.** (**A**) Overlay of X-ray crystal structures of human (red) and *C. elegans* (blue) START domains. Calculated RMSD was 1.3 Å over 103 atoms. (**B**) X-ray crystal structures of START domains from different STARD proteins.

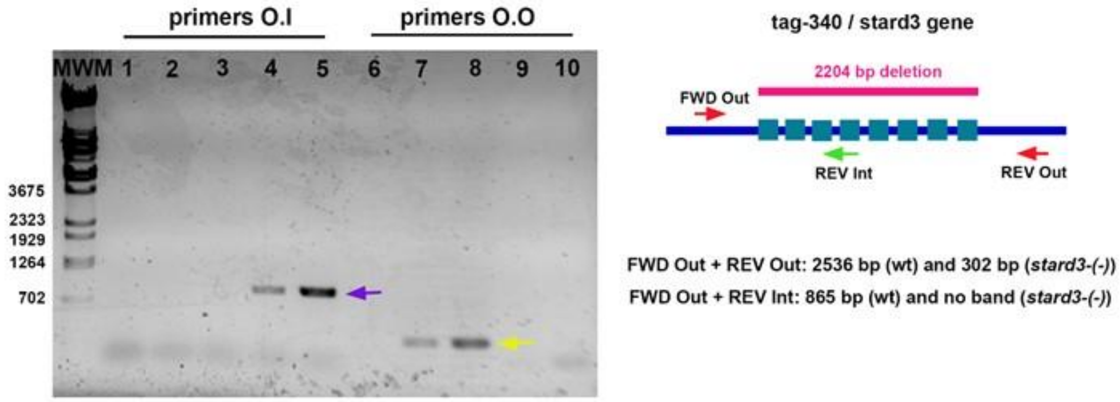


**Supplementary Figure 6: (A)** Agarose gel showing PCR amplification of the *stard3* sequence in wild type (N2) and the *stard3-(-)* mutant. Lanes 1 and 6: no DNA (negative control); 2 and 7: *stard3-(-)* DNA, 1/100 dilution; 3 and 8: *stard3-(-) DNA*, 1/10 dilution; 4 and 9: wild-type DNA, 1/100 dilution; 5 and 10: wild-type DNA, 1/10 dilution. Primers O.I: FWD Out + REV Int; primers O.O: FWD Out + REV Out. Molecular weight is expressed in base pairs. The purple and yellow arrows indicate the 865 and 302 bp bands, respectively. It was not possible to detect the 2536 bp band that should be amplified from Primers FWD Out and FWD Int in the wild type genotype due to the low efficiency of the standard PCR reaction. **(B)** Schematic representation of the *tag-340/stard3* gene in *C. elegans*. Light blue boxes represent exons. The genetic deletion in *stard3-(-)* is represented in magenta and the primers used for PCR confirmation are depicted as red and green arrows.

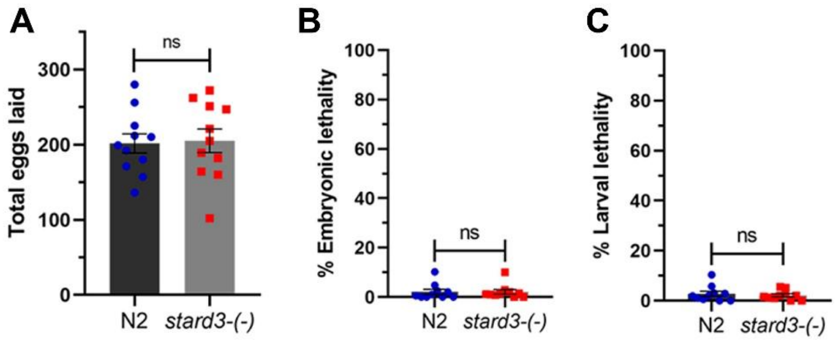


**Supplementary Figure 7.** Mean number of eggs laid **(A)**, percentage of embryonic lethality **(B)**, and larval lethality **(C)** are shown for N2 and *stard3-(-)* worms grown on standard NGM medium (supplemented with 13 µM of cholesterol) at 20ºC. A minimum of 10 P0 worms were analyzed for each condition. Error bars represent SEM.
